# Distinct effects of amyloid and tau deposition on eigenvector centrality during hippocampal down-regulation: a real-time fMRI virtual reality closed-loop neurofeedback study with CSF biomarkers

**DOI:** 10.1101/654426

**Authors:** Stavros Skouras, Jordi Torner, Patrik Andersson, Yury Koush, Carles Falcon, Carolina Minguillon, Karine Fauria, Francesc Alpiste, Kaj Blenow, Henrik Zetterberg, Juan D. Gispert, José L. Molinuevo, for the ALFA Study

## Abstract

Hippocampal down-regulation is associated with genetic predisposition to Alzheimer’s disease (AD), neurodevelopmental processes and disease symptoms. Resting state eigenvector centrality (EC) patterns resemble those of FDG-PET in AD, they can predict self-regulation performance and they are related to functional compensation across the pathophysiological continuum of AD. We acquired cerebrospinal fluid (CSF) biomarkers from a cognitively unimpaired sample at risk for AD (N=48), to investigate the effect of β- amyloid peptide 42 (Aβ42) and phosphorylated tau (p-Tau) levels on EC during the down-regulation of hippocampal subfield cornu ammonis 1, with real-time fMRI closed-loop neurofeedback. Controlling the effects of confounding variables (age, sex, number of *APOE* ε4 alleles, cognitive reserve, brain reserve and hippocampal down-regulation performance), CSF Aβ42 levels correlated positively with EC in the anterior cingulate cortex (BA24, BA32) and primary motor cortex (BA4). CSF p-Tau levels correlated with EC positively in the ACC (BA32, BA10) ventral striatum (caudate, nucleus accumbens, putamen) and left primary somatosensory cortex (BA2), as well as negatively in the posterior cingulate cortex, precuneus, cuneus and left frontal pole (BA9). Controlling for CSF biomarkers and other prognosis variables, age correlated negatively with EC in the midcingulate cortex, insula, primary somatosensory cortex (BA2) and inferior parietal lobule (BA40), as well as positively with EC in the inferior temporal gyri. Taken together, we identified patterns of functional connectomics in individuals at risk of AD during hippocampal down-regulation, which resemble those found during resting state at advanced AD stages. Moreover, we provide a standard paradigm to replicate and extend this work on a global level. This opens new avenues for further research applications, which quantify and monitor disease progression, by identifying early alterations in the self-regulation of brain function, with potential for non-invasive prognostic screening.

**Highlights:**

- ACC centrality decreases with early Aβ42
- ACC centrality increases with p-Tau
- PCC centrality decreases with p-Tau
- MCC centrality decreases in healthy aging

## Introduction

Alzheimer’s disease (AD) poses a global threat to millions of lives and the sustainability of public healthcare (Prince et al., 2015). Recent progress is shifting the mainstream dual clinic-pathological concept of AD towards a pathophysiological *continuum*, with progression being monitored *in vivo* through the study of cerebrospinal fluid (CSF) and positron emission tomography (PET) biomarkers (Jack et al., 2018). The study of the brain alterations associated with the presence of abnormal levels of biomarkers in unimpaired individuals may shed light on factors and mechanisms associated to cerebral resilience or vulnerability to early AD pathology.

Eigenvector centrality (EC) is one of the most advanced connectomic metrics (Bonacich, 1972; Langville and Meyer, 2006) that can consider global brain patterns of functional connectivity, in high image resolution and without *a priory* assumptions, to derive relative estimates of influence for nodes and clusters within networks (Borgatti, 2005, Lohmann et al., 2010; Wink et al., 2012). Previous studies have shown that EC reveals similar patterns to FDG-PET when comparing AD patients to healthy controls (Adriaanse et al., 2016), and that AD patients present significant differences in EC, in the anterior cingulate cortex (ACC), paracingulate gyrus, cuneus and occipital cortex (Binnewijzend et al., 2014). An independent study found that increased EC in the cingulate cortex and thalamus is related to compensatory mechanisms across the pathophysiological continuum of AD (Skouras et al., 2019a). However, all previous investigations of EC in AD have been limited to task-free (aka resting state) fMRI, despite evidence suggesting that functional tasks which engage specific brain networks affected by a disease, lead to stronger effects and consequently to higher discriminative power between patients and controls (Finn et al., 2017; Greene et al., 2018).

Across AD patient studies and healthy memory studies, the single brain region with the most important role has been established to be the hippocampus (Kim 2011; Schwindt and Black, 2009). Hippocampal subfield CA1 in particular, appears to be a region that is integral to contextual episodic memory (Penner and Mizumori, 2012; Mizumori et al., 1999; Leutgeb et al., 2004; Vazdarjanova and Guzowski, 2004). CA1 presents atrophy linked to AD pathology but no volume loss related to normal aging (Frisoni et al., 2008; Wilson et al., 2004; Yushkevich et al., 2015). Given that hippocampal hyperactivity appears to precede amyloid accumulation (Leal et al., 2017) and to be more prominent in subjects at genetic risk for AD (Tran et al., 2017), it is important to develop a standard framework for the investigation of hippocampal regulation abilities, that are conceptually related to neural reserve (Barulli and Stern, 2013; Stern et al., 2018). Real-time, closed-loop neurofeedback (NF) enables to study the self-regulation of specific brain regions during interactive functional tasks (Sitaram et al., 2017) and offers the possibility to be combined with virtual reality (VR) that promotes participants’ experimental compliance, through perceptual immersion in engaging and ecologically valid scenarios (Chirico et al., 2017; Krokos et al., 2018).

Here, we fused state-of-the-art, disruptive technologies (i.e. fully automated electrochemiluminescence immunoassay, real-time functional neuroimaging and VR), aiming to create an interactive and entertaining task to investigate the neural correlates of hippocampal down-regulation, associated with core AD CSF biomarkers and healthy aging, in a sample of cognitively unimpaired participants at risk for AD. We hypothesized that we may identify patterns of functional connectomics during hippocampal down-regulation that resemble those found during resting state at advanced AD stages. Specifically, we expected to find measurable differences of EC in the occipital cortex (Binnewijzend et al., 2014), the inferior parietal lobule, the thalamus and the cingulate cortex (Binnewijzend et al., 2014; Skouras et al., 2019a).

## Results

### Aβ42

Controlling for the effects of age, sex, number of *APOE* ε4 alleles, hippocampal volume, cognitive reserve and NF performance, CSF Aβ42 levels showed a significant positive correlation with EC in the ACC (BA24, BA32) and primary motor cortex (BA4); Fig. 2, Table 2. Note that Aβ42 levels in CSF are inversely proportional to the extent of β-amyloid plaque accumulation in the brain (Strozyk et al., 2003; Grothe et al., 2017); that is, abnormal Aβ42 biomarkers were related to low EC in the ACC and BA4.

**Fig. 1:**
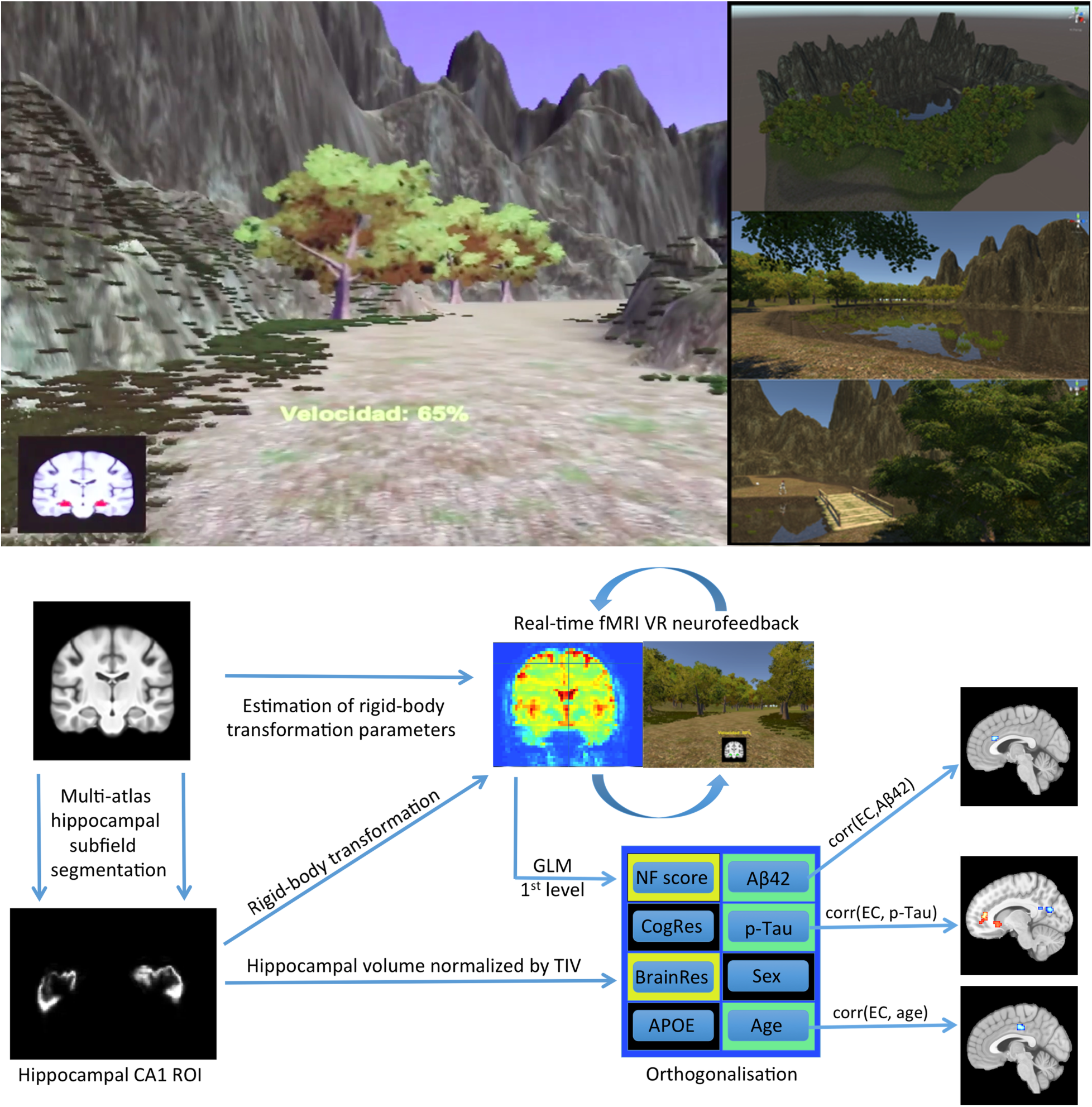
Virtual reality environment and neuroimaging pipeline. The VR NF paradigm developed for the study employed principles of passive, sensory-aided, operant conditioning and featured 570 NF signals per session. To maintain a balanced perceptual experience across participants, task difficulty adapted to individual performance dynamically, aiming to drive CA1 activity in each participant to the minimum possible. Using previously acquired anatomical images, multi-atlas hippocampal subfield segmentation localized and segmented hippocampal subfield CA1, prior to real-time scanning. With every real-time functional volume, moment-to-moment changes in hippocampal CA1 activation effected inverse changes of velocity in VR. Offline statistical modelling was used to derive a measure of NF regulation performance and to perform EC mapping (i.e. to estimate how much influence each brain region exerts during hippocampal CA1 down-regulation with NF). Abbreviations: fMRI ∼ functional magnetic resonance imaging; VR ∼ virtual reality; CA1 ∼ cornu ammonis 1; TIV ∼ total intracranial volume; GLM ∼ general linear model; NF ∼ neurofeedback; CogRes ∼ cognitive reserve; BrainRes ∼ brain reserve; APOE ∼ apolipoprotein genotype; EC ∼ eigenvector centrality; Aβ42 ∼ amyloid-beta 42; p-Tau ∼ phosphorylated tau.

**Figure 2.**
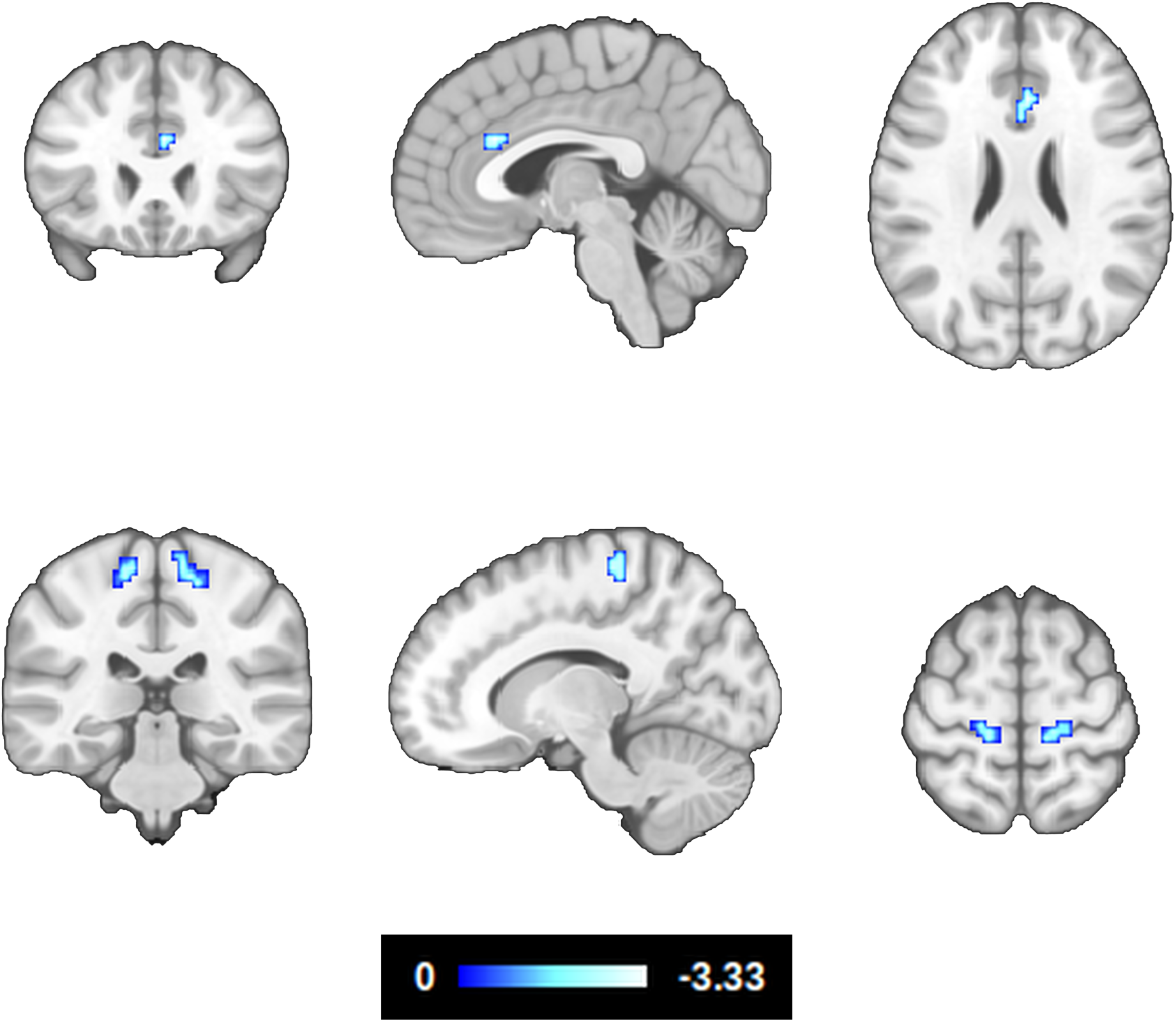
Correlation between eigenvector centrality during hippocampal down-regulation and β-amyloid deposition. The effects of age, sex, number of APOE-E4 alleles, hippocampal volume, cognitive reserve and NF performance, were modeled and controlled (z > 2.326, P < 0.05 whole-brain corrected). Note that CSF Aβ42 levels are inversely proportional to the extent of Aβ plaque deposition in the brain.

### p-Tau

Controlling for the effects of age, sex, number of *APOE* ε4 alleles, hippocampal volume, cognitive reserve and NF performance, p-Tau levels in CSF showed a significant positive correlation with EC in the ACC (BA32, BA10) ventral striatum (caudate, nucleus accumbens, putamen) and left primary somatosensory cortex (BA2). CSF p-Tau levels, also showed a significant negative correlation with EC in the PCC, PCu, Cu and left frontal pole (BA9); Fig. 3, Table 2.

**Figure 3.**
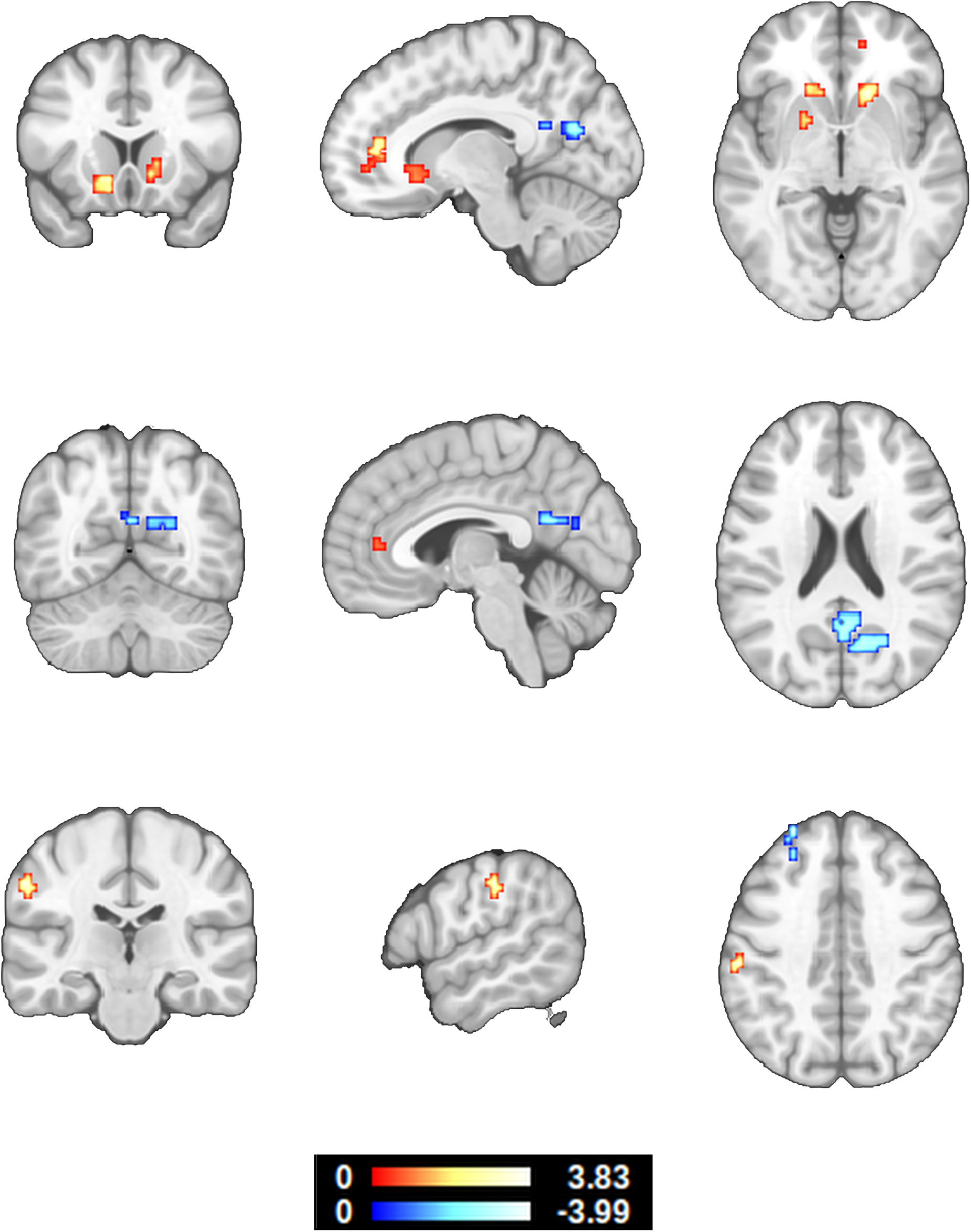
Correlation between eigenvector centrality during hippocampal down-regulation and CSF p-Tau levels. The effects of age, sex, number of APOE ε4 alleles, hippocampal volume, cognitive reserve and NF performance were modeled and controlled (z > 2.326, P < 0.05 whole-brain corrected). Note that CSF p-Tau levels are proportional to the presence of neurofibrillary tangles in the brain. These results suggest that AD-characteristic EC differences in the ACC may occur earlier than previously believed.

### Healthy Aging

Controlling for the effects of sex, number of *APOE* ε4 alleles, hippocampal volume, cognitive reserve, NF performance and the CSF p-Tau by Aβ42 ratio (Maddalena et al., 2003), age presented a significant negative correlation with EC in the midcingulate cortex (MCC), insula, primary somatosensory cortex (BA2) and inferior parietal lobule (BA40). Age also presented a significant positive correlation with EC in the inferior temporal gyri; Fig. 4, Table 2.

**Figure 4.**
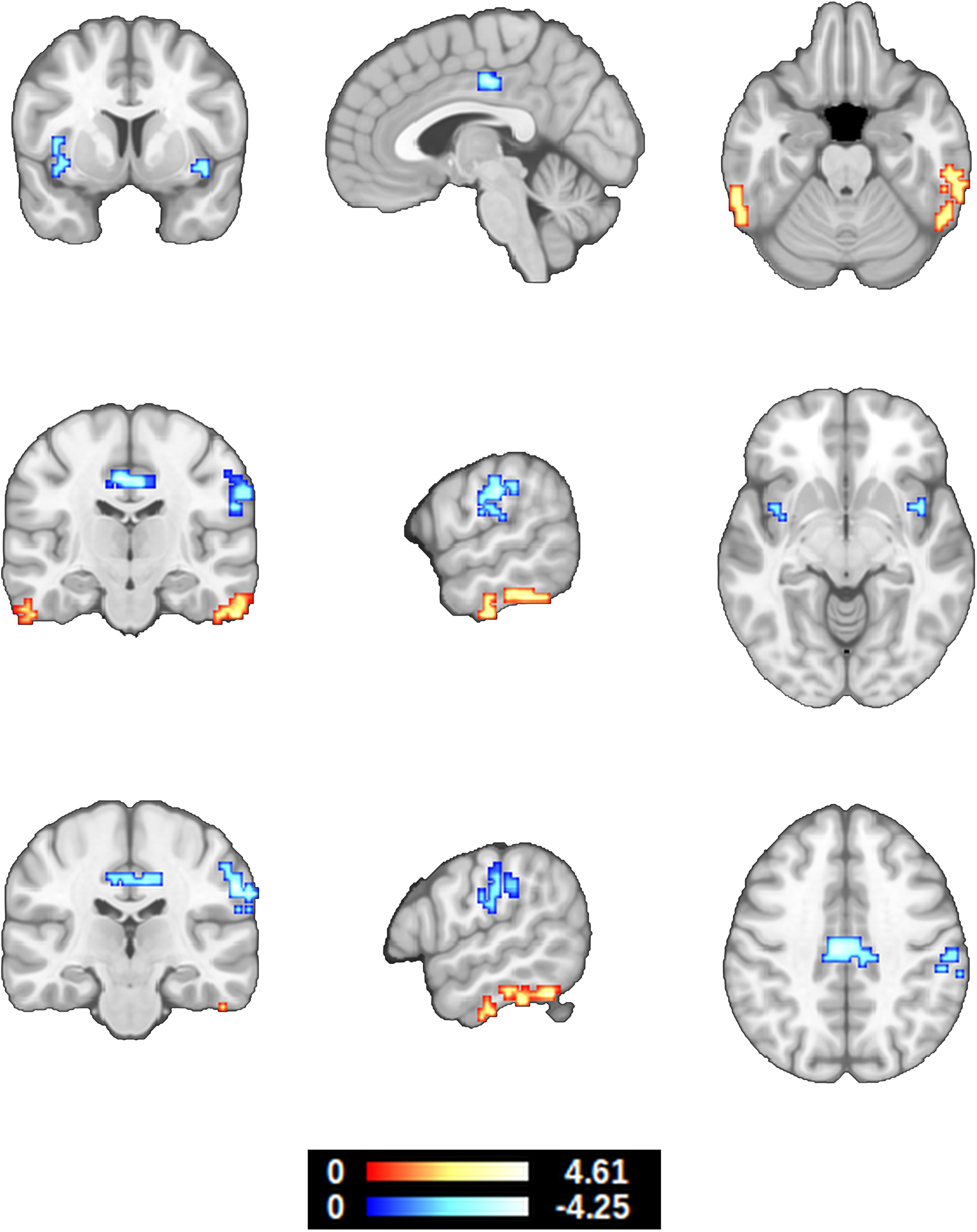
Correlation between eigenvector centrality during hippocampal down-regulation and age. The effects of sex, number of APOE ε4 alleles, hippocampal volume, cognitive reserve, NF performance, CSF Aβ42 and p-Tau levels were modeled and controlled (z > 2.326, P < 0.05 whole-brain corrected). In relation to Fig. 3, these results show that EC in the cingulate and BA2 present the opposite patterns in healthy aging.

## Discussion

Adding to our knowledge, this is the first real-time NF study utilizing CSF biomarker data. The results in cognitively unimpaired participants at risk for AD, corroborate that differences of EC in the cingulate cortex play an important role in the pathophysiological continuum of AD and show for the first time that such differences may begin at a very early pathophysiological stage. The latter finding suggests that hippocampal self-regulation tasks, enabling functional connectomic analyses, can be of benefit in revealing information of clinical relevance to AD progression.

During hippocampal down-regulation, the decreased EC in the ACC and primary motor cortex, that is associated with abnormally decreased CSF Aβ42 levels and, by extension, to increased Aβ plaque deposition in the brain, stands out as a potential neural correlate of elevated amyloid deposition; particularly because the ACC is among the first regions where amyloid deposition can be detected with PET imaging (Grothe et al., 2017). Due to the characteristics of our sample (Table 1), these findings represent early pathophysiological alterations, without objectively measurable impact on overall memory performance, that relate to the brain network involved in the regulation of hippocampal hyperactivity. They suggest that in subjects with abnormal Aβ42 deposition, the ACC is less influential in regulating hippocampal CA1 (adjusted for risk factors and task performance). According to current models, the ACC, together with the anterior insula, as well as parts of the prefrontal cortex, parietal lobule, ventral striatum and thalamus, comprise the brain network that is primarily responsible for learning to voluntarily regulate other brain areas through NF training (Sitaram et al., 2017). That provides a mechanistic explanation for the reason most of these areas, specifically, are engaged by the present VR NF paradigm, leading to enhanced sensitivity in relation to early alterations of their connectivity and systemic network function.

**TABLE 1:** Descriptive statistics and noncollinearity of measured variables. Abbreviations: FSCRT ∼ free and cued selective reminding test; CRQ ∼ cognitive reserve questionnaire; TIV ∼ total intracranial volume; NF ∼ neurofeedback; NC ∼ non-carriers; HT ∼ heterozygotes; HM ∼ homozygotes. Asterisks mark uncorrected significance at conventional alpha levels.

| | Descriptives | Age | Aβ42 | p-Tau | t-Tau | p-Tau/Aβ42 |
| --- | --- | --- | --- | --- | --- | --- |
| **Age** | M = 62.705<br>SD = 4.628 |  | R = 0.129<br>P = 0.382 | R = 0.369<br>P = 0.010** | R = 0.373<br>P = 0.009** | R = 0.224<br>P = 0.126 |
| **Aβ42** | M = 1186.200<br>SD = 424.678 | R = 0.129<br>P = 0.382 |  | R = 0.195<br>P = 0.184 | R = 0.276<br>P = 0.057 | R = -0.580<br>P < 0.001*** |
| **p-Tau** | M = 18.233<br>SD = 9.007 | R = 0.369<br>P = 0.010* | R = 0.195<br>P = 0.184 |  | R = 0.987<br>P < 0.001*** | R = 0.625<br>P < 0.001*** |
| **t-Tau** | M = 220.273<br>SD = 86.623 | R = 0.373<br>P = 0.009** | R = 0.276<br>P = 0.057 | R = 0.987<br>P < 0.001*** |  | R = 0.553<br>P < 0.001*** |
| **p-Tau / Aβ42** | M = 0.017<br>SD = 0.010 | R = 0.224<br>P = 0.126 | R = -0.580<br>P < 0.001*** | R = 0.625<br>P < 0.001*** | R = 0.553<br>P < 0.001*** |  |
| **FCSRT**<br>**Total recall** | M = 44.292<br>SD = 3.989 | R = -0.177<br>P = 0.227 | R = -0.202<br>P = 0.168 | R = 0.004<br>P = 0.976 | R = 0.014<br>P = 0.921 | R = 0.125<br>P = 0.398 |
| **FCSRT**<br>**Retention Index** | M = 0.980<br>SD = 0.066 | R = 0.088<br>P = 0.554 | R = 0.065<br>P = 0.658 | R = -0.046<br>P = 0.756 | R = -0.015<br>P = 0.921 | R = -0.083<br>P = 0.576 |
| **FCSRT**<br>**Delayed cued recall** | M = 3.708<br>SD = 1.675 | R = 0.250<br>P = 0.087 | R = -0.026<br>P = 0.861 | R = 0.143<br>P = 0.331 | R = 0.107<br>P = 0.468 | R = 0.156<br>P = 0.291 |
| **FCSRT**<br>**Delayed free recall** | M = 11.479<br>SD = 1.935 | R = -0.217<br>P = 0.138 | R = -0.087<br>P = 0.558 | R = -0.129<br>P = 0.383 | R = -0.090<br>P = 0.543 | R = -0.073<br>P = 0.621 |
| **FCSRT**<br>**Total free recall** | M = 28.562<br>SD = 5.379 | R = -0.307<br>P = 0.033 | R = 0.016<br>P = 0.913 | R = -0.128<br>P = 0.387 | R = -0.089<br>P = 0.547 | R = -0.137<br>P = 0.353 |
| **FCSRT**<br>**Total cued recall** | M = 15.729<br>SD = 4.409 | R = 0.239<br>P = 0.101 | R = -0.219<br>P = 0.134 | R = 0.175<br>P = 0.233 | R = 0.133<br>P = 0.369 | R = 0.305<br>P = 0.035* |
| **FCSRT**<br>**Total delayed recall** | M = 15.187<br>SD = 1.214 | R = 0.004<br>P = 0.978 | R = -0.197<br>P = 0.180 | R = -0.005<br>P = 0.974 | R = 0.008<br>P = 0.956 | R = 0.114<br>P = 0.439 |
| **CRQ score** | M = 16.479<br>SD = 3.724 | R = -0.012<br>P = 0.935 | R = 0.046<br>P = 0.755 | R = 0.013<br>P = 0.927 | R = 0.025<br>P = 0.868 | R = -0.081<br>P = 0.582 |
| **Hippocampal**<br>**volume / TIV** | M = 0.005<br>SD = 0.001 | R = -0.026<br>P = 0.859 | R = -0.012<br>P = 0.933 | R = 0.044<br>P = 0.764 | R = 0.068<br>P = 0.645 | R = -0.025<br>P = 0.864 |
| **Hippocampal down-**<br>**regulation NF score** | M = -0.038<br>SD = 0.462 | R = 0.219<br>P = 0.135 | R = 0.087<br>P = 0.557 | R = 0.215<br>P = 0.143 | R = 0.198<br>P = 0.176 | R = 0.115<br>P = 0.434 |
| **APOE ε4 allele**<br>**carrier status** | n<sub>NC</sub> = 19<br>n<sub>HT</sub> = 19<br>n<sub>HM</sub> = 10 | R = -0.699<br>P < 0.001*** | R = -0.325<br>P = 0.024* | R = -0.288<br>P = 0.047* | R = -0.290<br>P = 0.046* | R = 0.025<br>P = 0.864 |

**TABLE 2.** Results of eigenvector centrality mapping, in relation to CSF biomarkers and age, corrected for whole-brain multiple comparisons (P < 0.05) The outermost right column indicates the maximal z-value of voxels within a cluster (with the mean z-value of all voxels within a cluster in parentheses). CSF Aβ42 and p-Tau levels were orthogonalized to age, sex, number of APOE ε4 alleles, cognitive reserve, brain reserve and hippocampal down-regulation performance. For consistency, CSF Aβ42 values were multiplied by −1 during modelling, to align results and colormaps with the direction of the pathophysiological continuum of AD. Age was orthogonalized to the CSF p-Tau by Aβ42 ratio, sex, number of APOE ε4 alleles, cognitive reserve, brain reserve and hippocampal down-regulation performance. Abbreviations: MNI ∼ Montreal Neurological Institute; Aβ42 ∼ amyloid-beta 42; p-Tau ∼ phosphorylated tau; BA ∼ Brodmann Area; ACC ∼ Anterior Cingulate Cortex; MCC ∼ Midcingulate Cortex; PCC ∼ Posterior Cingulate Cortex; PCu ∼ Precuneus; Cu ∼ Cuneus; NAc ∼ Nucleus Accumbens.

| **B-amyloid correlation with Eigenvector Centrality during CA1 down-regulation** |  |  |  |
| --- | --- | --- | --- |
| **Brain regions** | **MNI**<br>**coordinates** | **Cluster size**<br>**(mm<sup>3</sup>)** | **z-value**<br>**max (mean)** |
| ACC, BA24, BA32 | 3 24 28 | 405 | -3.34 (-2.65) |
| L BA4 | -21 -27 70 | 405 | -2.73 (-2.49) |
| R BA4 | 18 -33 61 | 594 | -2.91 (-2.56) |
| **P-Tau correlation with Eigenvector Centrality during CA1 down-regulation** |  |  |  |
| **Brain regions** | **MNI**<br>**coordinates** | **Cluster size**<br>**(mm<sup>3</sup>)** | **z-value**<br>**max (mean)** |
| ACC, BA32 (BA10) | 12 42 13 | 513 | 3.31 ( 2.66) |
| R Caudate, NAc, Putamen | 15 21 -2 | 513 | 3.53 ( 2.86) |
| L Caudate, NAc, Putamen | -12 18 -5 | 756 | 3.84 ( 2.87) |
| L BA2 | -57 -24 43 | 297 | 3.54 ( 2.79) |
| PCC, PCu, R Cu | 21 -66 22 | 1674 | -3.36 (-2.69) |
| L Frontal pole, BA9 | -27 51 43 | 297 | -4.00 (-2.68) |
| **Age correlation with Eigenvector Centrality during CA1 down-regulation** |  |  |  |

| Brain regions | MNI<br>coordinates | Cluster size<br>(mm<sup>3</sup>) | z-value<br>max (mean) |
| --- | --- | --- | --- |
| R ITG | 57 -54 -23 | 1539 | 4.62 (3.01) |
| R ITG | 57 -21 -32 | 837 | 3.94 (2.83) |
| L ITG | -63 -45 -20 | 999 | 3.38 (2.77) |
| L ITG | -57 -18 -29 | 459 | 2.98 (2.65) |
| MCC, BA24, BA31 | -6 -21 43 | 1593 | -4.10 (-2.94) |
| R BA40, BA2 | 57 -24 37 | 2565 | -4.26 (-2.83) |
| L BA40, BA2 | -54 -30 28 | 675 | -3.92 (-2.89) |
| R Insula, BA13 | 42 6 -5 | 297 | -3.15 (-2.75) |
| L Insula, BA13 | -36 3 -2 | 648 | -3.35 (-2.60) |

With elevated CSF p-Tau levels, EC decreases in the prefrontal cortex and the PCC, while EC increases in the ACC and ventral striatum. That is, during early stages, Aβ42 accumulation in the brain and tau phosphorylation, seem to have opposite effects on EC in the ACC. The ACC is also involved in the brain signature of cognitive resilience, which is related to its metabolic capacity (Arenaza-Urquijo et al., 2019). The possibility of EC in the ACC initially decreasing and later on increasing, in early response to the subsequent steps of a pathophysiological cascade, resembles the concurrent pattern of default mode network (DMN) connectivity, that increases proportionally to CSF Aβ42 levels within the range of normal amyloid values, but decreases proportionally to CSF Aβ42 levels within the elevated range of abnormal amyloid accumulation (Palmqvist et al., 2017). This is supported by the decrease of EC in the PCC/PCu/Cu that comprises the main DMN hub (Smith et al., 2009; Spreng et al., 2010, Andrews-Hanna et al., 2010). Moreover, recent evidence suggests that baseline ECM in the PCC/PCu region is predictive of first-session DMN NF regulation learning (Skouras and Scharnowski, 2019). Overall, the EC differences associated with increased CSF p-Tau levels and by extension to the formation of neurofibrillary tangles in the brain, resemble results found in advanced AD patients with dementia, in the ACC and occipital cortex (Binnewijzend et al., 2014). In this context, the present results corroborate the most relevant literature and support shifting the timeframe for the detection of aberrant functional alterations, earlier than previously evidenced, to the presymptomatic stage of AD. Of particular importance, the present findings suggest that aberrant EC initially precedes neurodegeneration, rather than resulting from it. This is supported by complementary evidence showing minimal overlap between the ACC cluster and regions with gray matter reductions in AD patients (Binnewijzend et al., 2014).

Left BA8, the cluster presenting decreased EC in the left prefrontal cortex, is one of the main loci of autobiographical memory (Janata, 2009) and appears to be suitable as an accessible target region for non-invasive neurostimulation intervention studies (Reinhart and Nguyen, 2019). Following the interpretational framework proposed by recent work with resting-state EC and AD CSF biomarkers (Skouras et al., 2019a), given the involvement of the ACC, PFC and ventral striatum in the NF learning network (Sitaram et al., 2017), it is feasible that the decreasing EC of the PFC is being compensated by increasing EC in the ACC and ventral striatum; the latter of which is affected by amyloid deposition only at a relatively advanced stage (Grothe et al., 2017). Considering the crucial roles of the hippocampus, ventral striatum and cingulate in affective processing (Dalgleish, 2004), these findings support proposing that some of the earliest effects along the pathophysiological continuum of AD are related to dysfunctions of affective processing, similarly to almost all neuropsychiatric pathologies (Mannie et al. 2007; Hoekert et al. 2007; Connan et al. 2003; Leppännen et al. 2006; Kang et al. 2012). Additionally, in a mouse model of AD, selective neurodegeneration in the ventral tegmental area (VTA) at pre-plaque stages, resulted in lower dopamine outflow in the hippocampus and nucleus accumbens (NAc) and correlated with impairments of synaptic plasticity in CA1, as well as memory deficits and dysfunction of reward processing (Nobili et al., 2017). The VTA is particularly responsive to NF training (Macinnes et al., 2016) and outgoing fibers from the VTA connect directly to the hippocampus (Penner and Mizumori, 2012; Gasbarri et al., 1994). Thereby, it is also feasible that the increased EC in the dopaminergic ventral striatum compensates for early aberrancies in VTA function that are undetectable with 3T whole-brain fMRI. In addition, a contemporary model proposes that the hippocampus, the prefrontal cortex and the VTA, form the most crucial components of the long-term memory network (Penner and Mizumori, 2012).

As hypothesized, during hippocampal down-regulation, cognitively unimpaired subjects with elevated CSF p-Tau levels, exhibit a similar increase in EC, as the one present during resting state in AD patients (Binnewijzend et al., 2014), despite important improvements in normalization methods across studies (Klein et al., 2009; Avants et al., 2008). This corroborates a replicable resting-state finding that was previously only measurable when investigating across control subjects and patients with dementia due to AD (Binnewijzend et al., 2014; Skouras et al., 2019a). This observation suggests that the developed VR NF task may indeed be hypersensitive to preclinical alterations in brain function. In contrast, an important effect of healthy aging is decreased EC in the cingulate cortex, during hippocampal down-regulation. This corroborates previous evidence from task-free fMRI, showing that healthy older adults (age M=63, SD=7) present significantly lower EC in the cingulate than healthy younger adults (age M=24, SD=4) (Antonenko et al., 2018). Overall, the effect of healthy aging on EC across the cingulate cortex (Fig. 4) is opposite to that of elevated CSF p-Tau levels (Fig. 3), clinical symptoms (Binnewijzend et al., 2014), and AD progression (Skouras et al., 2019a). Thereby, current evidence suggest that the cingulate decreases in centrality in healthy aging and increases in centrality in AD, with noticeable differences even in a very early phase. The effect of healthy aging on EC in the ITG, points to the possibility that in healthy aging, increasing resources are being devoted to semantic memory (Binney et al., 2010). To our knowledge, there is no previous evidence of EC correlation with age in the ITG, however no previous ECM aging study controlled for the effect of any CSF biomarkers. Moreover, recent evidence suggests cognitive decline may emerge from functional decoupling within a neural circuit composed of temporal and frontal regions, that is integral to monitoring real-world information and storing it into memory (Reinhart and Nguyen, 2019).

In general, the EC findings could be different using a hippocampal up-regulation task, because up-regulation and down-regulation learning can be negatively correlated (Skouras and Scharnowski, 2019). It is important to determine the specificity of the present findings to hippocampal down-regulation tasks and to replicate them in longitudinal studies that include both down-regulation and up-regulation, as well as sham NF conditions. It is equally important to validate the tentative interpretations offered here, using the same self-regulation paradigm in studies involving patients with mild cognitive impairment (MCI). With these aims, we have made the developed VR environment publicly available as open-source software, to enable replication studies and multi-center investigations within a standard framework (see section *Code Availability*). The latter might enable the aggregation of sufficient datasets to derive accurate functional neuroimaging-based diagnostic models using machine learning algorithms.

We have demonstrated that amyloid deposition results in aberrancy with regards to the functional brain network utilized to down-regulate hippocampal subfield CA1. Further, significant differences in EC associated with CSF biomarkers in clinical AD, are also measurable in presymptomatic stages. Moreover, we provide a standard paradigm to replicate and extend this work on a global level. This opens new avenues for further research applications, which quantify and monitor disease progression, by identifying early alterations in the self-regulation of brain function, with potential for non-invasive prognostic screening.

## Methods

### Participants

Participants comprised of 48 adult volunteers (age M = 62.705 years; SD = 4.628) from the ALFA (Alzheimer’s and Families) project (Molinuevo et al. 2016), many of them descendants of AD patients and *APOE ε4* allele carriers, hence presenting increased risk for AD. All participants were highly functional and without neurological or psychiatric history at the time of scanning. 22 participants presented CSF Aβ42 levels < 1098, 15 participants presented CSF p-Tau levels > 19.2 and 7 of those participants met both criteria. Within the preceding six months, participants had completed the Free and Cued Selective Reminding Test (FCSRT) (Grober et al., 2009); Table 1. Participants had previously also completed the Cognitive Reserve Questionnaire (CRQ); a questionnaire comprised of 10 questions whose total score serves as a proxy for cognitive reserve (Rami et al., 2011). Thorough quality control was applied in advance, to exclude datasets having more than 10% invalid functional volumes due to movement and acquisitions during which technological complications, fatigue, discomfort or sleep had occurred. The local ethics committee ‘CEIC-Parc de Salut Mar’ reviewed and approved the study protocol and informed consent form, in accordance with current legislation.

### CSF sampling

CSF was collected by lumbar puncture between 9 and 12 a.m. in polypropylene tubes. Samples were processed within 1 hour and centrifuged at 4°C for 10 minutes at 2000 g, stored in polypropylene tubes and frozen at −80°C. Core AD CSF biomarkers (namely Aβ42 and p-Tau) were determined using cobas Elecsys® assays (Hansson et al., 2018).

### Apolipoprotein E genotyping

Proteinase K digestion and subsequent alcohol precipitation, was utilized to obtain DNA from the blood cellular fraction. Samples were genotyped for two single nucleotide polymorphisms (rs429358 and rs7412) and the number of *APOE* ε4 alleles was determined for each participant (Molinuevo et al., 2016). Results are displayed in Table 1.

### Image acquisition

All scanning was performed in a single 3T Philips Ingenia CX MRI scanner (2015 model). Pre-NF scanning comprised of a 3D T1-weighted sequence of 240 sagittal slices with Voxel Resolution = 0.75 × 0.75 × 0.75 mm3, Repetition Time (TR) = 9.90 ms, Echo Time (TE) = 4.60 ms, Flip Angle = 8; and a diffusion-weighted sequence of 66 axial slices with Voxel Resolution = 2.05 × 2.05 × 2.20 mm3, TR = 9000 ms, TE = 90 ms, Flip Angle = 90, featuring 72 non-collinear directions (b = 1300 s/mm2) and 1 nongradient volume (b = 0 s/mm2). During NF, echo planar imaging was used with Voxel Resolution = 3× 3× 3 mm3, TR = 3000 ms, TE = 35 ms, Matrix Size = 80 × 80 voxels, Field of View = 240 mm and interleaved slice acquisition with an interslice gap of 0.2 mm (45 slices, whole brain coverage).

### Structural image processing

The standard FreeSurfer pipeline was applied on T1-weighted images, to estimate each participant’s normalized hippocampal brain volume, as a proxy of brain reserve, by dividing total hippocampal volume by total intracranial volume (Cavedo et al., 2012). Using the Advanced Normalization Tools (Avants et al., 2009) (ANTs v2.x; RRID: SCR_004757), the N4 nonparametric non-uniform intensity normalization bias correction function (Tustison et al., 2010; Tustison and Avants, 2013) was applied on all T1 images, followed by an optimized blockwise non-local means denoising filter (Coupé et al., 2008). Multi-atlas segmentation with joint label fusion (Wang et al., 2013) segmented hippocampal subfields to derive probabilistic maps for CA1, which were thresholded at P = 0.9 to create the NF target ROI masks.

### VR paradigm and real-time NF

The design of the VR paradigm was guided by the following objectives: a) we used VR to make the task immersive, engaging and entertaining; b) that in turn enabled making the task particularly long, 30 minutes, to maximize 1^st^-level statistical power; c) we employed sliding-window closed-loop NF to achieve an optimal design for analysis of functional connectomics; d) we narrowed the NF target ROI to hippocampal subfield CA1, that presents atrophy in AD but not in healthy aging (Frisoni et al., 2008; Wilson et al., 2004; Yushkevich et al., 2015), while also being consistently implicated in memory encoding (Kim, 2011) and differing in activity in patients (Schwindt and Black, 2009). Prior to the acquisition of CSF biomarkers, the task was validated in a larger, partly independent sample (77% overlap), by showing that hippocampal down-regulation was associated with genetic predisposition to AD, neurodevelopmental processes and bilateral cohesion of hippocampal function (Skouras et al., 2019b).

The VR environment and real-time computations have been described in detail elsewhere (Torner et al., 2018). Briefly, the VR environment had been developed in the game-engine Unity (Unity Technologies ApS, San Francisco, CA, US) (Fig. 1). During a 30-minute-long scanning session, participants were immersed in the VR environment and could run in a fixed, circular path. The experimental task was to explore different mental strategies, aiming to achieve the maximum velocity and to traverse the maximum distance possible, while at the same time attending to features of the VR environment, remembering them and considering whether they changed from round to round. The first 30 functional volumes served to establish a baseline of hippocampal activity, for each participant. Subsequently, with every TR, the most recent shift in hippocampal CA1 activity was compared to reference measures of change derived from the preceding 90 seconds. A 5% decrease of VR velocity was triggered by increases in hippocampal activity and a 5% increase of velocity was triggered by decreases in hippocampal activity. Every 30 volumes, the velocity was reset to 50%. At any given moment, the current velocity was displayed as a percentage of the maximum velocity possible and a green or red signal was superimposed on a coronal brain view, reflecting the direction of the most recent change (Fig. 1). Real-time image preprocessing consisted of rigid-body registration to a reference volume from the same scanning session, temporal high-pass filtering at a cut-off frequency of 1/200 Hz (Tarvainen et al., 2002), as well as voxel efficiency weighting (Stoeckel et al., 2014) via voxel-wise normalization within the 90-second sliding time-window (for extensive details please refer to Torner et al., 2018).

### Functional image processing

Offline functional image processing consisted of standard preprocessing using SPM12 (Statistical Parametric Mapping, RRID:SCR_007037) and the MIT connectivity toolbox (Connectivity Toolbox, RRID:SCR_009550), including slice-time correction, realignment and reslicing of functional volumes, denoising via regression of average white matter timeseries, average CSF timeseries, 24 Volterra expansion movement parameters and scan-nulling regressors (Lemieux et al., 2007) produced by the Artifact Detection Tools (ART; RRID: SCR_005994). Functional data and CA1 ROIs were normalized to the ICBM MNI template, featuring sharp resolution and detailed gyrification, as well as high signal-to-noise ratio with minimum bias (Fonov et al., 2019), using SyGN (Tustison and Avants, 2013; Avants et al., 2011) and a custom template derived from the same population (Skouras et al., 2019a). Temporal highpass filtering with a cut-off frequency of 1/90 Hz and spatial smoothing using a 3D Gaussian kernel and a filter size of 6 mm at FWHM, were performed using the Leipzig Image Processing and Statistical Inference Algorithms (LIPSIA v2.2.7 (2011), RRID:SCR_009595; Lohmann et al., 2000). Finally, EC was computed voxel-wise similarly to a previous study with AD patients (Skouras et al., 2019a).

### Statistical Analysis

Pearson’s r was used as a metric of similarity in first-level GLM, to quantify NF performance per participant. A linear vector coding continuous down-regulation, as the target performance for NF, was compared to NF moment-to-moment regulation, measured by the realigned average CA1 timeseries, to produce a measure of NF performance for each participant. First, we investigated whether the unique variance associated with CSF Aβ42 levels was reflected in EC during the NF task. For consistency, CSF Aβ42 values were multiplied by −1 during modelling, to align results and colormaps with the direction of the pathophysiological continuum of AD. CSF Aβ42 values were orthogonalised to potential confounding variables; specifically sex, age, number of *APOE* ε4 alleles, hippocampal volume, cognitive reserve and NF performance. This resulted in an orthogonal 2^nd^-level design matrix that was used for GLM of the unique effect of Aβ42 on EC during VR NF. Whole-brain functional connectivity maps were corrected for multiple comparisons using 10,000 iterations of Monte Carlo simulations, with a pre-threshold of Z > 2.326 (P < 0.01) and a corrected significance level of P < 0.05. Similarly, we investigated for a possible correlation between p-Tau and EC, controlling for the same confounding variables. Finally, we investigated the effect of healthy aging on EC during VR NF, by orthogonalizing age to all potential confounding factors, including the CSF p-Tau by Aβ42 ratio (Maddalena et al., 2003). All scale variables were normalized prior to orthogonalization via the recursive Gram-Schmidt orthogonalisation of SPM12 (SPM, RRID:SCR_007037). For quality control, we confirmed that each variable of interest was completely orthogonal to all covariates in its respective design matrix, as well as to all original vector data, while remaining in high correlation with its own original vector. This ensured investigating meaningful observations and effects due to the unique variance in each variable of interest.

### Code and Data accessibility

The VR environment is publicly available as open-source software via the ‘VR_multipurpose_v1.0’ repository (GitHub; RRID:SCR_002630). Data used and software developed for the analysis can been made available to researchers for non-commercial purposes, following agreement and approval by the Barcelonaβeta Brain Research Center’s Data and Publications Committee.

## Acknowledgements

This work has received funding from the European Union’s Horizon 2020 research and innovation programme under the Marie Sklodowska-Curie action grant agreement No 707730.

This publication is part of the ALFA (ALzheimer and FAmilies) study which receives support from la Caixa Foundation (LCF/PR/GN17/10300004). The authors would like to express their most sincere gratitude to the ALFA project participants, without whom this research would have not been possible. Authors would like to thank Roche Diagnostics International Ltd. for kindly providing the kits for CSF core AD biomarker analyses.

Collaborators of the ALFA study are: Jordi Camí, Anna Brugulat-Serrat, Raffaele Cacciaglia, Marta Crous-Bou, Carme Deulofeu, Ruth Dominguez, Carolina Herrero, Carles Falcon, Xavi Gotsens, Nina Gramunt, Oriol Grau-Rivera, Laura Hernandez, Gema Huesa, Jordi Huguet, María León, Paula Marne, Tania Menchón, Marta Milà-Alomà, Grégory Operto, Maria Pascual, Albina Polo, Sandra Pradas, Aleix Sala-Vila, Gemma Salvadó, Gonzalo Sánchez-Benavides, Sabrina Segundo, Anna Soteras, Marc Suárez-Calvet, Laia Tenas, Marc Vilanova and Natalia Vilor-Tejedor.

